# Perturbation-Aware Neural ODE (pNODE) Learns Microbiome Dynamics from Clinical Data and Predicts Gut-Borne Bloodstream Infections in Patients Receiving Cancer Treatment

**DOI:** 10.1101/2025.11.26.690798

**Authors:** Isaac Stamper, Brendan Aeria, Joao Xavier

## Abstract

Disruption of the gut microbiota during cancer treatment, particularly in allogeneic hematopoietic cell transplantation (allo-HCT), contributes to adverse clinical outcomes, including gut-borne bloodstream infections. Accurately forecasting microbial population dynamics under clinical perturbations, such as antibiotic administration, could inform treatment strategies to reduce infection risk. However, traditional models like the Generalized Lotka-Volterra (gLV), which consider only pairwise interactions with constant sign and magnitude, are limited in capturing the real-world nonlinear dynamics of multispecies microbiomes following ecosystem disturbances. Here, we introduce a perturbation-augmented Neural Ordinary Differential Equation (pNODE) framework that flexibly models microbial population dynamics in continuous time, integrating both microbial abundances and time-resolved antibiotic perturbations. Using synthetic and real clinical data from over 1,000 allo-HCT patients, we show that pNODEs outperform gLV in predictive accuracy, robustness to noise, and forecasting critical events, such as the intestinal expansion of an opportunistic pathogen. Notably, we demonstrate that running a pre-trained pNODE in generative mode to simulate prospective *Enterococcus* abundance trajectories from an initial sample and antibiotic timeline yields scores that accurately predict subsequent bloodstream infections in held-out cohorts, outperforming baseline and ground-truth-based predictors. Our findings demonstrate the potential of pNODEs as a next-generation tool for modeling clinical microbiome dynamics, with applications for predicting infections in immunocompromised patients hospitalized for cancer treatment.

## Introduction

Cancer treatment regimens, particularly allogeneic hematopoietic cell transplantation (allo-HCT), can disrupt the gut microbiota with implications for patient outcomes [1–5]. Chemotherapy and radiation suppress the immune system, leaving patients vulnerable to life-threatening infections and necessitating broad-spectrum antibiotic treatment [6, 7]. These antibiotics cause collateral damage to the gut microbiota, reducing its biodiversity and the numbers of commensal strict anaerobic bacteria and impairing key microbiome functions, such as resistance to expansion by opportunistic pathogens. As a result, opportunistic facultative anaerobes such as *Enterococcus faecium, Escherichia coli*, and *Klebsiella pneumoniae*, which are normally present at very low fractions (<1%), can expand to comprise >30% of the gut bacterial population, a phenomenon we call intestinal domination [1]. Intestinal domination facilitates translocation of these facultative anaerobes into the bloodstream, an aerobic environment, leading to systemic infections associated with higher morbidity and mortality [1, 8–10].

Despite clear evidence linking antibiotics, intestinal domination, and bloodstream infection, antibiotics are still administered with little consideration for their ecological impact [11]. This creates a self-reinforcing cycle: antibiotics damage the microbiota, increasing susceptibility to infection, which then prompts further antibiotic use and worsens the global problem of rising antibiotic resistance [12]. A new paradigm is needed in which antibiotics are selected based on their ecological impact on a patient’s gut microbiome.

A key step toward this much-needed paradigm shift is developing predictive models that forecast microbiome dynamics in response to clinical perturbations. Such models could inform treatment decisions that minimize disruption to the microbial community while reducing infection risk. However, reliably predicting microbial population dynamics remains a major challenge [13, 14]. Classical ecological models, such as the Generalized Lotka-Volterra (gLV), make simplifying assumptions that limit their applicability: they assume linear, pairwise, and time-invariant interactions in a homogeneous environment [15]. While gLV models have yielded interpretable insights in some settings [16–18], they are constrained in capturing the nonlinear, context-dependent dynamics of the intestinal microbiome, especially under complex perturbations such as antibiotics [19, 20].

To address these limitations, we propose expanding on Neural Ordinary Differential Equations (NODEs) as a flexible, data-driven alternative [21]. Unlike models with fixed functional forms, NODEs use neural networks to parameterize the system of differential equations governing microbial dynamics. This lets them learn arbitrary, nonlinear relationships directly from time-series data while operating in continuous time [22–24]. Recent work has explored related ODE-based formalisms in microbiome research [25–27]. Neural Jump ODEs (NJODEs) have been applied to detect antibiotic-associated anomalies in irregularly sampled infant microbiome trajectories [28]. NJODEs focus on estimating conditional mean and variance trajectories for anomaly scoring, rather than modeling ecological dynamics or forecasting the future composition of specific taxa. In contrast, our approach develops a perturbation-aware NODE framework that explicitly incorporates real antibiotic exposures and learns population-level microbial dynamics directly from clinical patient data. By operating on actual taxonomic abundances and treatment timelines, our method is designed for predictive ecological modeling in allo-HCT patients, where sampling is sparse, perturbations are strong, and clinical outcomes such as pathogen domination and bloodstream infection are central.

We call pNODE the expanded NODE framework modeling external perturbations. We evaluate its performance using both synthetic and clinical data. For synthetic validation, we conducted in silico experiments with known ground truth and evaluated pNODE’s ability to capture population dynamics, benchmarking it against the gLV model. For the clinical evaluation, we leveraged a longitudinal dataset of over 10,000 microbiota samples from more than 1,000 allo-HCT patients, including 16S rRNA sequencing, detailed antibiotic records, and clinical metadata [29]. We then applied a pre-trained pNODE in generative mode and demonstrated its potential to forecast microbial composition and to produce a score that predicts bloodstream infections caused by *Enterococcus* in prospectively analyzed patients receiving allo-HCT. We also used the model to visualize antibiotic-dependent flow fields in our previously developed TaxUMAP atlas (Schluter et al. 2023), suggesting that different antibiotic regimens drive microbial communities toward distinct ecological states, including *Enterococcus*-dominated attractors. Our results establish pNODEs as a promising tool for predictive modeling in clinical microbiome research and lay the groundwork for microbiome-aware, data-driven treatment strategies.

## Methods

### Clinical cohort and microbiota data

We analyzed a previously published longitudinal microbiome dataset of patients undergoing allogeneic hematopoietic cell transplantation (allo-HCT) at Memorial Sloan Kettering Cancer Center (MSK) [29]. This resource combines serial fecal microbiota profiles with detailed hospital records, including antibiotic administration, laboratory values, and clinical outcomes. We restricted our analysis to patients with at least two fecal microbiota samples and corresponding antibiotic data available in the time interval of interest. For a subset of samples, absolute bacterial load by 16S rRNA gene quantitative PCR (qPCR), providing estimates of total bacterial density per gram of stool.

### Taxonomic aggregation: class-level and genus-level state representations

Because the pNODE state vector *x*(*t*) can be defined at any taxonomic resolution, we constructed two representations and trained a separate pNODE at each scale. This allowed us to assess how taxonomic resolution influences learned microbiome dynamics and downstream prediction of clinical endpoints.

To reduce dimensionality, we aggregated taxa at the bacterial class level. Class-level abundances were obtained by summing within-class relative abundances, with very-low-abundance classes merged into a single “low-abundance” group. Because the *Bacilli* class in this cohort is clinically dominated by *Enterococcus faecium*, the pathogen most strongly linked to gut-borne bloodstream infections, we split *Bacilli* into a dedicated *Enterococcus* coordinate (genus-level) and a *Bacilli*_*not*_*Enterococcus* remainder so the predictor of clinical interest is resolved explicitly. After this aggregation, each microbiota sample is represented as a 14-dimensional vector *x* ∈ ℝ^14^ comprising 11 named classes plus the low-abundance group and the two Bacilli-derived coordinates.

For the genus-level representation, we retained genera with prevalence ≥5% and mean relative abundance ≥0.01% across the cohort; the remaining genera were grouped into per-class “removed” buckets so that no abundance is discarded. This yielded *N*_*genus*_ = 136 coordinates: 124 named genera passing the filter, 11 per-class “removed” buckets, and one “unassigned” bucket for reads not mapped to any genus.

Samples with missing or zero total counts after aggregation were excluded from modeling. For analyses requiring absolute abundances (gLV inference), we restricted to the subset of samples for which qPCR measurements were available and combined class-level relative abundances with total bacterial load to estimate class-specific absolute abundances.

### Antibiotic administration data and perturbation encoding

We extracted systemic antibacterial treatments from the hospital medication records curated in the “hospitalome” component of the dataset [29]. We retained only systemic antibiotics and excluded non-antibacterial agents (e.g., antivirals, antifungals) for the main analyses. We mapped individual antibiotic drugs to 15 antibiotic categories (e.g., carbapenems, fluoroquinolones, anti-pseudomonal β-lactams). For each patient, this produced a time-resolved record of antibiotic exposure intervals.

We encoded antibiotic treatment as a time-dependent perturbation function *p*(*t*). For each patient, *p*(*t*) is a 15-dimensional vector, where the *i*-th coordinate *p*_*i*_ (*t*) equals 1 if the patient received an antibiotic in category *i* at time *t* and 0 otherwise. This defines a function:

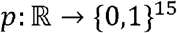

which may be generalized to continuous-valued perturbations *p*: ℝ ⟶, ℝ^15^ (e.g., antibiotic doses administered or actual drug concentrations measured in feces). We aligned antibiotic start and stop times to a continuous time axis measured in days relative to the nearest transplant date.

### Bloodstream infection outcomes

We obtained bloodstream infection (BSI) data from curated infection tables linked to the same cohort [29]. We focused on BSIs caused by *Enterococcus* spp., as these are strongly linked to high intestinal *Enterococcus* abundance (intestinal dominations) and have a clear ecological interpretation. For each patient, we identified the time of the first *Enterococcus* BSI and used this to define prediction windows for infection-risk models.

### Synthetic ecosystem simulations with a perturbation-augmented gLV model

To evaluate pNODE under controlled conditions with known ground truth, we constructed a synthetic 5-species microbial ecosystem governed by a generalized Lotka-Volterra (gLV) model with a single external perturbation. To mimic clinical data, we subsampled the dense trajectories at irregular time points and added Gaussian noise. We varied sampling density (observations per trajectory), the number of training trajectories, and observation noise level across a grid; for each combination, we simulated multiple independent ecosystems split into training and test sets.

### Generalized Lotka-Volterra inference

We fit a perturbation-augmented gLV model by regularized linear regression on finite-difference time derivatives, following Stein et al. (2013):

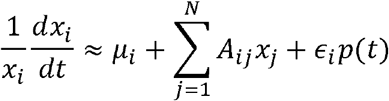

yielding estimates of growth rates *µ*, interactions *A*, and antibiotic-response coefficients *ϵ*.

For the clinical data, we restricted gLV inference to the subset of samples with qPCR-based absolute abundances. We applied the same regression at the class level to log-transformed absolute abundances and antibiotic exposures.

### Perturbation-aware Neural ODE (pNODE) architecture

The pNODE adopts a compositional formulation so that integrated outputs are valid relative-abundance vectors at every time. An unconstrained latent state *z*(*t*) ∈ ℝ^*N*^ is integrated by the network, and the observation-space state is read off as *x*(*t*_0_ = *so ftmax*(*z*(*t*)). The latent dynamics are governed by

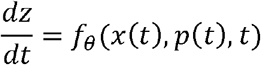

where *θ* denotes the neural network parameters and *p*(*t*) is the time-dependent antibiotic encoding, and *t* is the absolute time – days relative to transplant – concatenated to the input.

The function *f*_*θ*_ is a feed-forward network with SiLU activations (Linear(*N* + *P* + 1,64) ⟶ SiLU ⟶ Linear(64,32) ⟶ SiLU ⟶ Linear(32,*N*)); initial conditions are mapped as *z*(0) = log (max(*x*(0), 10^-6^)) and integrated using the Euler solver from torchdiffeq [21].

For the qPCR-restricted comparison against gLV, the pNODE is trained on the 14-dimensional class-level state described above. The genus-level pNODE (pNODEGenus) used for the full-cohort analysis uses the same architecture with wider hidden layers (128 and 64 instead of 64 and 32) for the higher-dimensional state. Each configuration is evaluated in its own state space: the full-cohort genus-level model is scored on the 136-dimensional genus vector, and the qPCR-subset class-level model on the 14-dimensional class vector.

### Training pNODE on synthetic ecosystems

For the synthetic gLV experiments, we trained pNODE to reconstruct full time-course dynamics from noisy, irregularly sampled data. For each simulated trajectory, the first observation served as the initial condition and the antibiotic schedule *p*(*t*) as input; the ODE solver integrated the pNODE forward to each observation time. The loss was mean absolute error (MAE) between predicted and ground-truth relative abundances, averaged over species and observation times in the batch.

We trained over a grid of dataset conditions (sampling density, noise level, number of trajectories). For each condition, we trained both models on the same training split and evaluated them on held-out trajectories with zero noise and regular sampling, scoring both on relative-abundance MAE. The pNODE’s predictions lie on the simplex by construction; we clipped the gLV’s predictions at zero and row-normalized them before scoring.

### Training pNODE on clinical microbiome data

For clinical data, we trained two pNODE configurations to make short-horizon forecasts at different taxonomic resolutions. We restricted the training objective to 7-day prediction windows in both cases, since this is the horizon on which follow-up samples are most consistently available per index sample; we then probed the model’s ability to extrapolate beyond this window at evaluation.

The training procedure was identical for both configurations. Each training example consisted of:

1. An “index” microbiota sample at time *t*_0_, represented as a state vector of relative abundances at the chosen taxonomic resolution.
2. The antibiotic perturbation function *p*(*t*) for the patient, defined for *t* ≥ *t*_0_.
3. All microbiota samples collected from the same patient within 7 days of *t*_0_.

For a given index sample, the ODE solver integrated the pNODE from *t*_0_ to each follow-up sample time *t*_*k*_ ∈ [*t*_0_,*t*_0_ + 7], using the patient-specific antibiotic function *p*(*t*) as input to *f*_*θ*_. At each follow-up time, we computed the MAE between the predicted and observed relative abundances. The loss for that index sample was the mean of these errors across all follow-up times and state dimensions. Mini-batches consisted of groups of index samples drawn from multiple patients. We excluded samples without any follow-up within 7 days from training.

The two configurations differed in their state representation and patient cohort. The class-level pNODE used the 14-dimensional class-level state and was trained on the qPCR-restricted subset of patients (for whom absolute abundances were available), enabling a direct comparison with gLV. The *genus-level* pNODE used the *N*_*genus*_-dimensional state and was trained on the full patient cohort, without a gLV comparison (which requires absolute abundances).

We partitioned patients into disjoint train/validation/test splits (60/20/20 %) so each sample appeared in only one split. For each clinical pipeline, we trained 20 independent replicates differing only in PyTorch’s random initialization seed, retained the parameter set with the lowest validation MAE per replicate, and reported all clinical metrics as the mean ± standard deviation across these 20 replicates. We assessed final predictive performance on the held-out test set by computing the MAE between predicted and observed relative abundances across all classes and time points.

### Comparing pNODE and gLV on clinical data

To compare pNODE with gLV on clinical data under fair conditions, we restricted the analysis to samples with qPCR-based absolute abundances, as required for gLV modeling. For each such sample, we constructed two representations:

- an absolute abundance vector for gLV inference, and
- a relative abundance vector for pNODE.

We trained a class-level gLV model on the absolute abundances and the pNODE on the corresponding relative abundances, using only data from the training patients. We then ran both models in generative mode on held-out test patients: starting from an initial sample, we integrated each model forward 7 days using the patient-specific antibiotic record. For each sample with a follow-up within 7 days, we scored predictions against observed abundances by MAE on relative abundances (gLV predictions clipped at zero and row-normalized). A “constant composition” baseline (microbiota fixed at the initial sample) was also evaluated, and the prediction horizon was varied to produce MAE-vs-window curves.

### Prediction of *Enterococcus* bloodstream infections from generative pNODE trajectories

To test whether a pre-trained pNODE could predict clinically relevant infections, we used the model in generative mode to predict microbiome composition itmeseries, which we then used to compute risk scores for *Enterococcus* BSIs. Critically, the pNODE used in this analysis was trained only to predict microbiota dynamics and was never trained directly on infection outcomes.

For each patient, each index sample, and each chosen prediction window length *w* (in days), we:

1. Took the class-level relative abundances at the index sample as the initial condition.
2. Supplied the patient-specific antibiotic perturbation function *p*(*t*) to the pNODE.
3. Integrated the pNODE forward from the index time to *t*_0_+ *w*.

From the resulting predicted trajectory, we extracted the *Enterococcus* coordinate and computed its mean over the *w*-day window. We used this scalar as a pNODE-based risk score for *Enterococcus* BSI occurrence in that window. We defined the binary outcome for each index sample and window as whether a documented *Enterococcus* BSI occurred within *w* days after the index sample (and not earlier).

We compared pNODE-based scores with two baselines: (i) the *Enterococcus* relative abundance at the index sample, and (ii) an interpolated ground-truth oracle, which builds a piecewise-linear interpolant through the index sample and all observed *Enterococcus* values within (*t*_0_,*t*_0_ + *w*] (clamping past the last sample), evaluates it on a 1-day grid, and takes the time-average. The oracle uses future measurements and is an upper bound for any *Enterococcus*-trajectory predictor. For each predictor and W, we computed ROC curves and AUCs on a held-out test set; AUCs are reported as mean ± SD across 20 pNODE replicates, with deterministic baselines contributing a single value.

### Visualizing perturbation-induced dynamics as flow fields in TaxUMAP space

To visualize the community dynamics learned by the genus-level pNODE, we projected microbiota compositions onto a two-dimensional TaxUMAP embedding and rendered the pNODE-induced dynamics as a flow field on that map [4]. We used the published TaxUMAP coordinates as a fixed reference space; because a precomputed-metric UMAP admits no out-of-sample transform, we embedded each pNODE-generated composition by out-of-sample projection, computed a combined genus- and family-level Bray-Curtis distance to the reference samples, and took the distance-weighted mean of its nearest reference coordinates. For this analysis, we used a pNODE trained without the explicit time input, so that the right-hand side is time-homogeneous given the antibiotic schedule, and the velocity field over composition space does not depend on a choice of day relative to transplant; forecast accuracy is unchanged (30-day MAE 0.061 vs 0.062 on the qPCR subset). To construct the flow field under a given antibiotic condition, we drew a large set of observed community states as seeds and advanced each one a short step (Δ*t* = 0.5 day) with the trained pNODE under that condition, using the perturbation function *p*(*t*); a no-antibiotic schedule provided the unperturbed control, and setting a single category active provided that antibiotic’s field. We embedded each state and its stepped endpoint, took the map-space displacement divided by Δ*t* as an instantaneous velocity, interpolated these velocities onto a regular grid by distance-weighted *k*-nearest-neighbor regression, and drew streamlines (masking grid regions with little support from real samples). We repeated this for each of the 12 most frequent perturbation states in the training data—the exact set of antibiotic categories co-active over a sample transition—one of which is the no-antibiotic control (14.5% of transitions); together, these 12 states cover 70.6% of training transitions. To indicate where the dynamics ultimately drive communities, we additionally integrated a diverse set of initial states to a long horizon (30 days) under each condition and overlaid the density of their endpoints, colored by dominant bacterial class, on the same map; we note that long-horizon endpoints extrapolate beyond the training window and reflect the learned model’s attractors rather than directly observed dynamics.

### Use of large language models

We used large language model-based assistants (Anthropic Claude, Grammarly and Microsoft Codex) in two capacities. First, to polish and edit manuscript text—proofreading, clarifying phrasing, and tightening prose drafted by the authors—but not to generate any section of the manuscript from scratch. Second, to assist in writing, debugging, refactoring, and documenting the analysis code (data preprocessing, model training, evaluation, and figure-generation scripts are all available in the public repository linked below). All scientific content, analyses, results, and interpretations are those of the authors, and the authors take full responsibility for the contents of this work.

### Code availability

Analyses used Python with NumPy, pandas, SciPy, scikit-learn, PyTorch, and torchdiffeq. Code is available at:

https://github.com/stampei1/Perturbation_NODE_microbiome_dynamics

The repository provides a single-command entry point (`python code/run.py demo|insilico|clinical|figures|all`) and SLURM submit scripts for HPC use; see `code/README.md&` and `CODE_OUTLINE.md` for details.

### Interactive web application

A browser-based explorer for the genus-level pNODE is available at: https://aeriab.github.io/pNODE_explorer/ where the forecast for any patient in the cohort can be recomputed under a user-edited antibiotic timeline and viewed as both a composition trajectory and a path on the TaxUMAP map of Fig. 5. The model is served as JSON and integrated client-side, so the site is fully static.

## Results

### Augmenting NODEs to account for extrinsic perturbations enables modeling microbiota dynamics during antibiotic treatment

The original NODE framework, which uses a neural network to model the time derivative of a multidimensional state variable, is well suited to describing system dynamics in the absence of external perturbations [21]. Here, we modified the architecture to model microbiome dynamics in the presence of extrinsic perturbations applied over fixed time intervals, a framework we call perturbation-aware NODE: (pNODE). We represent each patient’s antibiotic history as a time-resolved function *p*: ℝ ⟶ {0,1}^*P*^ with *p*_*i*_(*t*) = 1 when antibiotic *i* is administered at time *t*, and 0 otherwise. *p*(*t*) need not align with the irregular microbiota sampling times. To include perturbations, we augment the NODE function to depend on *p*(*t*) as well as *x*(*t*).

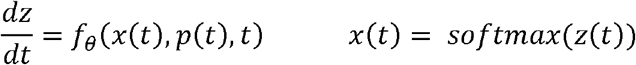

To implement this architecture, we add input nodes to the neural network for each dimension of the perturbation, so *f*_*θ*_ takes in vectors of length *N*+ *P* + 1, rather than just *N* — concatenating the current state *x*, the perturbation *p*(*t*), and the absolute time *t* (Fig. 1A).

**Figure 1.**
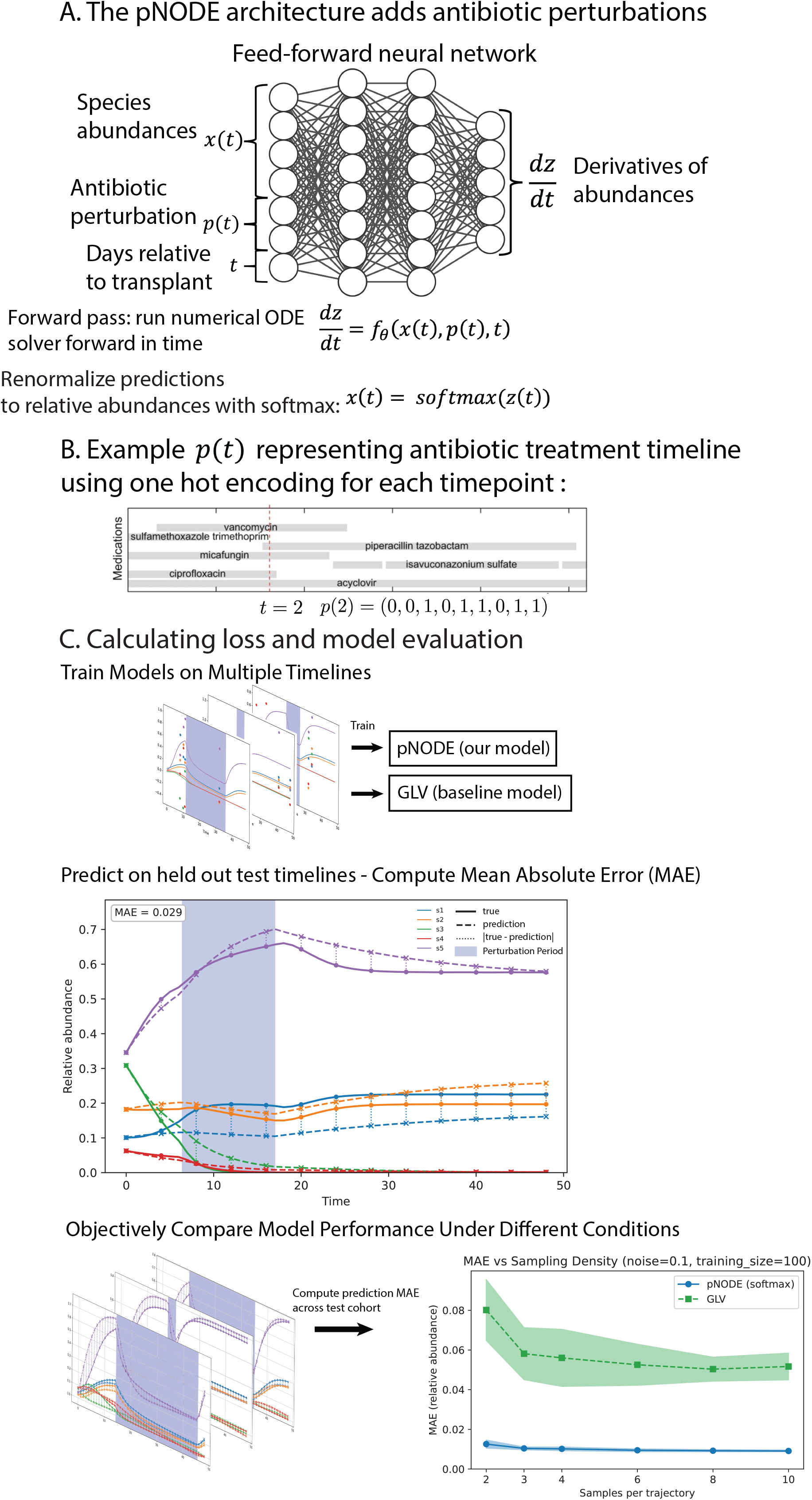
The perturbation-aware Neural ODE (pNODE) architecture. (A) Schematic of the pNODE architecture. A feed-forward neural network *f*_*θ*_ takes as input the current species abundances *x*(*t*), a time-resolved antibiotic perturbation vector *p*(*t*), and the days relative to HCT (time *t*), and outputs the instantaneous derivative 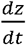, which a numerical ODE solver integrates forward in time. To make predictions, the numerical ODE solution is renormalized to relative abundances using a softmax: *x*(*t*) = *softmax*(z(*t*)) (B) Example of a patient antibiotic-treatment timeline encoded as *p*(*t*): at every integer time *t*, the active antibiotic categories are read off the medication record and concatenated into a one-hot binary vector. (C) Training and evaluation workflow. The pNODE and a Generalized Lotka-Volterra (gLV) baseline are trained on the same set of sampled trajectories, run in generative mode on held-out test trajectories to obtain predicted abundances at every sampling time, and scored by mean absolute error (MAE) against the true trajectories. Comparing the two models across a grid of training conditions yields the systematic benchmark presented in Figure 2 (illustrated for one slice: MAE vs sampling density at fixed noise = 0.1 and training-size = 100).

The pNODE is a special case of a Neural Controlled Differential Equations model [30]. In generative mode, *p*(*t*) is retrieved from the clinical record at time *t* and concatenated with *x*(*t*) before being fed into *f*_*θ*_ (Fig. 1B).

### pNODE outperforms gLV inference even in gLV-generated data

To validate pNODE, we constructed a synthetic 5-species ecosystem (N = 5) with one toggled antibiotic perturbation (P = 1), governed by a perturbation-augmented gLV model:

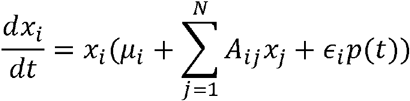

Here *x*_*i* is the abundance of species *i, µ*_*i*_ its intrinsic growth rate, *A* the pairwise interaction matrix (interpretable as the adjacency matrix of a directed ecological network), and ∈_*i*_ the antibiotic susceptibility of species i; *p*(*t*) is the binary perturbation (Fig. 2A).

**Figure 2.**
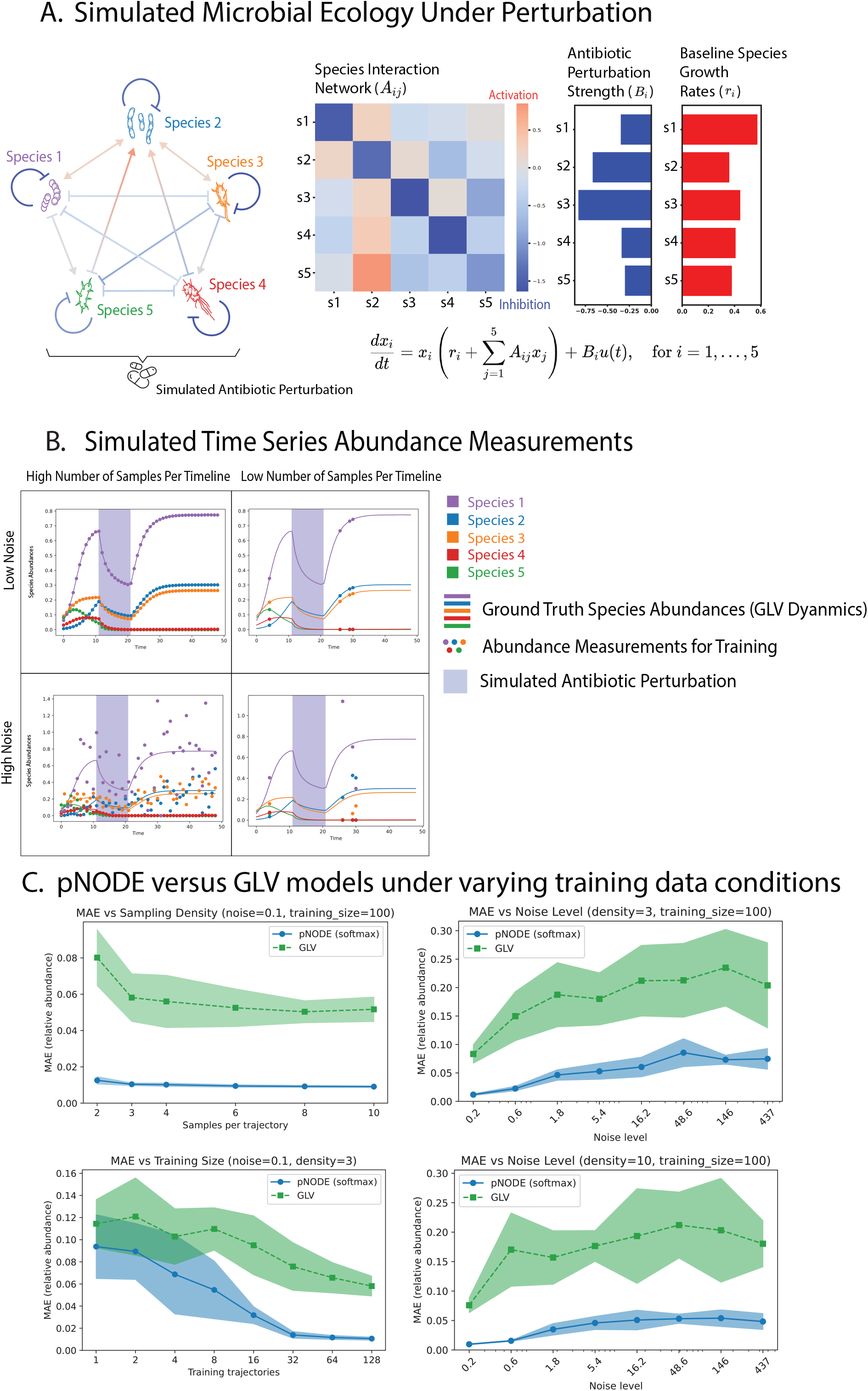
Synthetic benchmark: pNODE outperforms gLV inference under realistic clinical-data conditions. (A) Ground-truth 5-species ecosystem used to generate synthetic data. A perturbation-augmented gLV model is specified by a directed species-interaction network and its interaction matrix *A*_*ij*_ (red = activation, blue = inhibition), per-species antibiotic susceptibility coefficients *B*_*i*_, and intrinsic growth rates *r*_*i*_ . (B) Example simulated time series under four combinations of sampling density (high vs low) and observation noise (low vs high); solid curves are the gLV ground truth and circles are the noisy, sparsely sampled observations used to train the models. The shaded interval marks the simulated antibiotic-perturbation window. (C) Held-out test MAE of pNODE (blue) vs gLV (green) across the (noise, density, training-size) grid. Top-left: MAE vs sampling density at noise = 0.1, training-size = 100. Bottom-left: MAE vs training-cohort size at noise = 0.1, density = 3. Top-right and bottom-right: MAE vs noise level at fixed density = 3 and density = 10, respectively (training-size = 100). pNODE achieves lower MAE than gLV across the entire grid; the gap is largest under sparse, noisy, and data-poor conditions. Curves are mean across replicates; shaded bands are ± 1 SD. Full MAE-ratio heatmaps across the grid are provided in Supplementary Fig. S1.

We sampled ground-truth interaction parameters from a previously published multivariate-normal prior [18], generated trajectories from Latin-hypercube initial conditions with an adaptive Runge-Kutta solver, and applied an antibiotic perturbation over a randomly chosen time window for each trajectory. This yields a controlled synthetic benchmark with known ground-truth dynamics.

We then compared pNODE’s efficacy for learning these gLV ecosystems under perturbations with traditional gLV-based inference [16]. We compared how the models performed under realistic conditions observed in clinical data by varying the noise level and using irregular, sparse time sampling of the synthetic data used to train the models (Fig. 2B). Across >2,000 experiments, we trained both models on matched data while varying observation noise, training-set size (number of timelines), and the density at which the 48-point timelines were subsampled to mimic sparse clinical sampling.

One might expect that because a gLV model generates the ground truth, gLV inference should outperform pNODE under most conditions. Surprisingly, the opposite was true: across the (noise, density, training-size) grid we tested, the pNODE consistently achieved lower test-set MAE than the gLV (Fig. 2C; full MAE-ratio heatmaps in Supplementary Fig. S1). The pNODE’s advantage was largest in sparse, noisy, data-poor regimes that resemble real clinical microbiome data, but it persisted even under the densest sampling, lowest noise, and largest training-cohort conditions, where one would expect parametric gLV inference to converge to the true dynamics. These synthetic-data experiments showed that pNODEs can model complex ecosystems under external perturbations, even when classical models match the structure that generated the ground truth.

### Training pNODE to learn microbiota ecological dynamics from clinical data

We then trained pNODE on real-world clinical data using relative abundance measurements and antibiotic treatment timelines from the dataset described above. As detailed in Methods, we trained two configurations: a class-level pNODE (N = 14) on the qPCR-restricted subset (enabling comparison with a gLV baseline), and a genus-level pNODE (*N*_*genus*_ = 136) on the full patient cohort, both using the same 15-dimensional binary antibiotic vector (*P*= 15).

Once trained, we can run a pNODE in the generative mode to predict microbiome dynamics in response to a given treatment regimen (Fig. 3A,B). Generative-mode prediction requires only an initial fecal sample and the planned antibiotic regimen: starting from *x*(*t*_0_), we integrate *f*_*θ*_ forward in time using the patient’s *p*(*t*).We use the same solver during training, where observed follow-up samples are compared with predictions via a loss function, and gradients are propagated back through the solver to update *θ* [21]. The training algorithm iterates through index samples drawn from multiple patients, and mini-batches consist of groups of such samples (each yielding a 7-day prediction window; see Methods).

**Figure 3.**
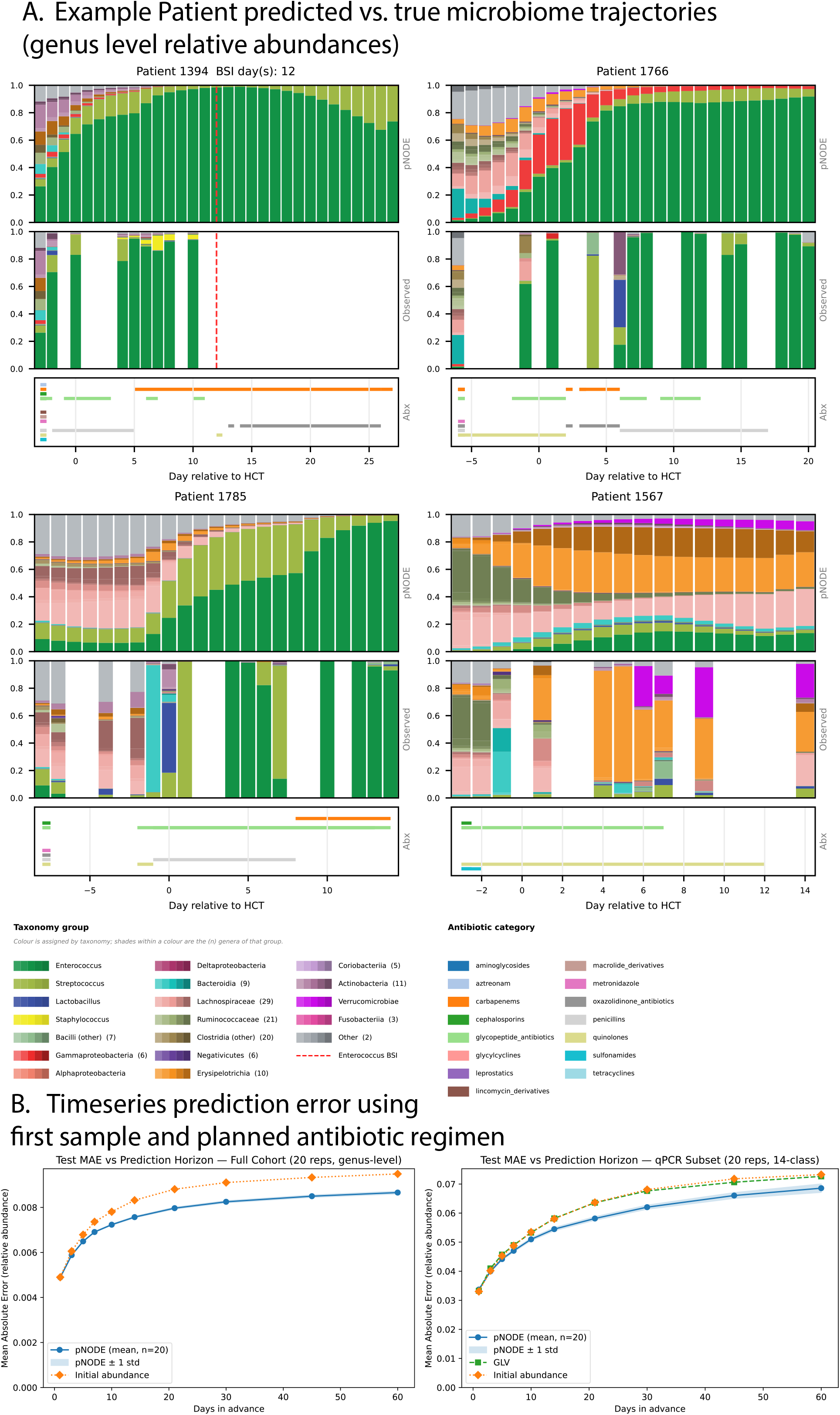
pNODE accurately predicts short-term microbiome dynamics in clinical data. (A) Four held-out patients (1394, 1766, 1785 and 1567) showing, for each patient, stacked genus-level relative abundances predicted by the full-cohort pNODE (top) and the observed abundances (middle), together with the antibiotic-exposure strip supplied to the model (bottom strip). Each trajectory is generated from that patient’s first sample in the window and their recorded antibiotic timeline alone — no later observation is used. Color is assigned by taxonomy rather than abundance: every genus is mapped to a palette group keyed on its class or family, and the genera within a group are given evenly spaced shades of that group’s anchor colour, so a clade reads as one colour family (legend, below the panels; the number after a group name is how many genera share that colour). Taxa are stacked in palette-group order so same-coloured relatives are adjacent, with *Enterococcus* pinned to the bottom of every stack. The vertical red dashed line marks a documented Enterococcus bloodstream-infection event (patient 1394, day 12). The pNODE tracks the major ecological transitions during antibiotic exposure and recovery, including both progression to *Enterococcus* domination (1394, 1766, 1785) and communities that remain diverse (1567). Predicted timelines for additional patients, along with an interactive version of the antibiotic strip that lets you edit the regimen to explore counterfactual exposures, are available in the web explorer at https://aeriab.github.io/pNODE_explorer/. (B) Test MAE vs prediction horizon for the full-cohort genus-level pNODE (left; pNODE vs initial-abundance baseline) and the qPCR-subset class-level pNODE (right; pNODE vs gLV vs initial-abundance baseline). Points are the mean across 20 pNODE replicates and shaded bands are ± 1 SD; gLV and initial-abundance baselines are deterministic. pNODE achieves the lowest MAE across the horizon range in both pipelines, from 3 days ahead in the qPCR subset and across the entire 1-60-day range in the full cohort.

We assessed the pNODE by generating new trajectories and comparing with held-out data. For the full-cohort analysis, we used the genus-level pNODE described in Methods, which operates at higher feature resolution and is scored directly on the genus-level state. To propagate training variability, we trained 20 independent replicates differing only in random initialization seed and report all clinical metrics as mean ± standard deviation across replicates. The genus-level pNODE captured a substantial portion of the population dynamics in the patient data, predicting relative abundances over the following week with 7-day-ahead MAE 0.007 on held-out full-cohort patients (Fig. 3A).

For a fair comparison with gLV, which requires absolute abundances, we restricted this analysis to the qPCR subset of samples [29]. We trained both a gLV and pNODE model on this subset, using absolute abundance measurements to train the gLV and the original relative abundance measurements to train the pNODE. We then compared their performance by running both models in generative mode to predict microbiota population dynamics over the next 7 days on held-out data. We also compared with a naive model that assumes that all subsequent time points equal the abundances of the initial sample (the microbiota population is constant in time).

The results showed that pNODE achieves 7-day MAE ≈*0*.*047* ± 0.0004, while the gLV achieves MAE ≈ *0*.*049* and the constant-composition baseline ≈ 0.049 (all on held-out data). The pNODE’s advantage widens at longer horizons: at 30 days ahead the pNODE achieves MAE *0*.*062* ± *0*.*001* vs gLV *0*.*068* and constant *0*.*068*, and at 60 days ahead *0*.*069* ± *0*.*002* vs *0*.*073* and *0*.*073*. When we changed the test prediction window size, the pNODE continued to outperform both the gLV and the constant-composition baseline (Fig. 3B). A complementary analysis on the held-out cohort also showed that the pNODE outperforms the gLV at predicting the direction (i.e., sign of change) of *Enterococcus* over the next 1-60 days (Supplementary Fig. S2). Together, these results support the pNODE for building predictive models of clinical events that occur due to shifts in the microbiomta composition, such as the occurrence of a gut-borne infection that is normally preceded by an intestinal expansion.

### Running pre-trained pNODE in generative mode generates time series that predict patients with bloodstream infections

We next tested the pNODE-generated trajectories to predict bloodstream infections, a task for which it had not been trained. Consistent with the ecological premise of the analysis, the measured *Enterococcus* relative abundance in fecal samples rises sharply in the days preceding a documented *Enterococcus* BSI relative to time-matched non-BSI controls (Fig. 4A). For each prediction window of length W, we integrated the pre-trained pNODE from an initial sample under the patient’s antibiotic timeline and used the mean predicted *Enterococcus* relative abundance over W days as a risk score, scoring it against the binary outcome of an *Enterococcus* BSI within W days via the area under the ROC curve (AUC). We compared the generative pNODE score against three alternatives (Methods): (i) initial *Enterococcus* relative abundance at the index sample (the simplest clinical baseline); (ii) the analogous gLV generative prediction on the qPCR subset (mean predicted *Enterococcus* over the window, clipped at zero and row-normalised); and (iii) an interpolated ground-truth oracle using observed *Enterococcus* values within the window, serving as an upper bound for any *Enterococcus*-trajectory-based predictor.

**Figure 4.**
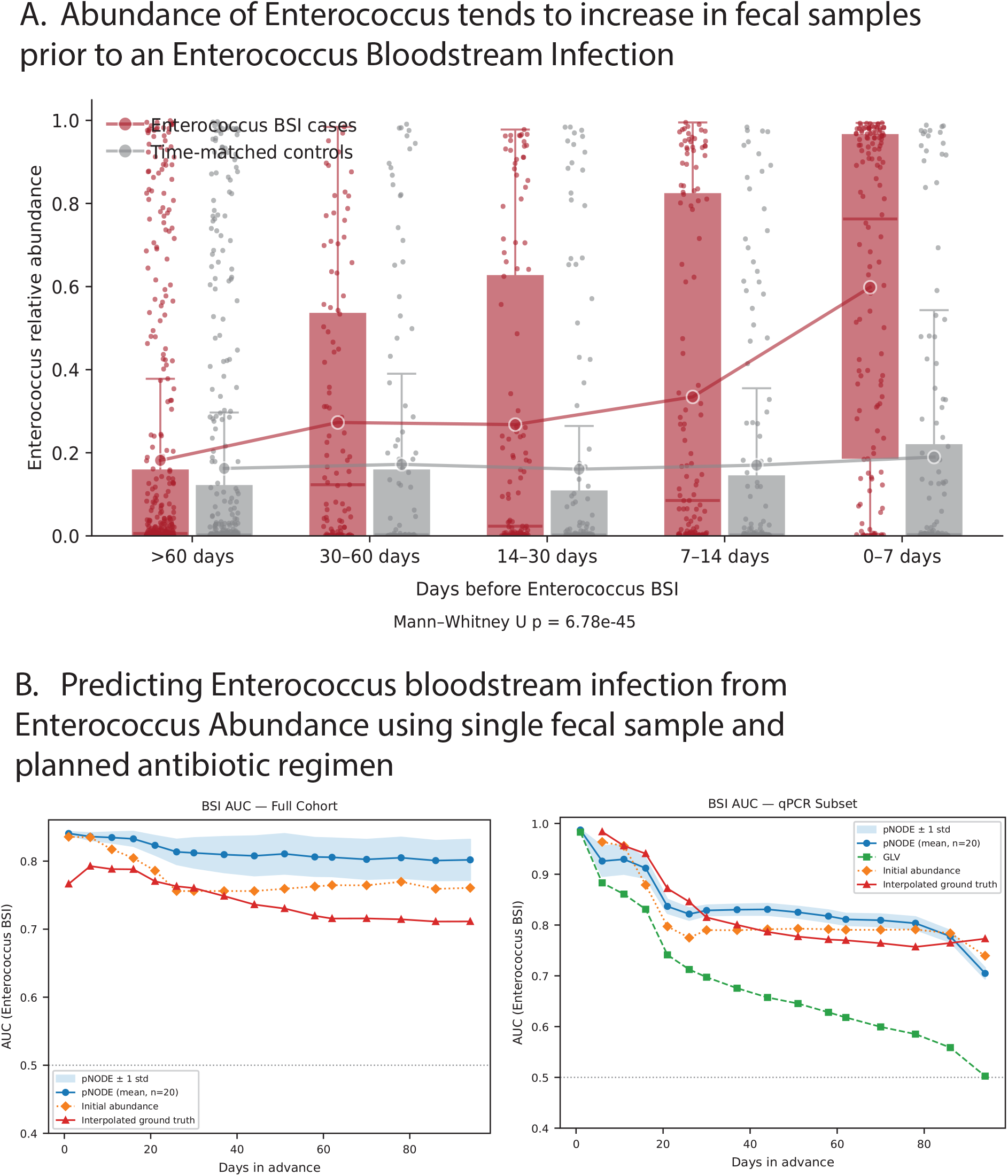
Generative-mode pNODE predictions identify patients at risk for *Enterococcus* bloodstream infection. (A) Time-matched case-control comparison of measured *Enterococcus* relative abundance in held-out fecal samples binned by days before an *Enterococcus* BSI event. Cases (red) are samples from patients who went on to develop an *Enterococcus* BSI; controls (grey) are time-matched samples from HCT-day-matched non-BSI patients. Boxplots show median and interquartile range with whiskers at 1.5×IQR; individual samples are overlaid as semi-transparent dots, and the bold lines connect the means across bins. The case-control difference is highly significant across the time series (Mann-Whitney U test, p = 6.78e-45). *Enterococcus* abundance rises markedly in cases in the 0-14 days preceding the BSI event, while controls remain low. (B) AUC for *Enterococcus* BSI prediction as a function of the prediction horizon W (days). Left: full-cohort genus-level pNODE (blue) vs initial-*Enterococcus* baseline (orange) vs interpolated ground-truth oracle (red). Right: qPCR-subset class-level pNODE (blue), gLV-generative (green), initial-*Enterococcus* baseline (orange), and interpolated ground-truth oracle (red). pNODE points are the mean across 20 replicates with ± 1 SD shaded; other curves are deterministic. The pNODE matches or exceeds the oracle at mid- and long-range horizons in the full cohort, and outperforms gLV in the qPCR subset across all horizons. The grey dashed line marks the chance level (AUC = 0.5).

We used these procedures on the same test dataset and computed AUC values for different prediction window sizes. On the qPCR subset, the pNODE-predicted trajectories outperformed gLV-predicted trajectories across the full range of horizons tested, exceeded the initial-*Enterococcus* baseline from W = 16 to W = 78 days, and matched or exceeded the interpolated ground-truth oracle at mid- and long-range horizons; for example, at W = 30 days, pNODE AUC = 0.83 ± 0.01 vs gLV 0.70, initial 0.79, and interpolated GT 0.82, and at W = 58 days pNODE 0.82 ± 0.02 vs gLV 0.63, initial 0.79, and interpolated GT 0.77. On the full-cohort genus-level pNODE, trained on 713 patients rather than 156, the gap to the oracle is more pronounced: pNODE remains at AUC ≈0.81–0.82 across W = 30–58 days while the interpolated ground-truth declines from 0.76 to 0.72 over the same range. Notably, the pNODE’s advantage widens with larger window sizes, dramatically outperforming the comparators at long horizons (Fig. 4B). A time-matched case-control comparison of the measured *Enterococcus* prior to BSI events further supports the underlying signal pNODE encodes: cases show progressively higher Enterococcus abundance as the BSI event approaches, relative to time-matched non-BSI controls (Fig. 4A).

### The pNODE encodes an antibiotic-dependent flow field over community-composition space

Because the pNODE defines a continuous vector field *f*_*θ*_ over composition space, we can inspect the learned dynamics directly. We projected the microbiota onto a two-dimensional TaxUMAP map, in which each point is a community and nearby points have similar composition, and rendered the pNODE-predicted motion as streamlines on this map (Fig. 5A,B; see *Methods*). Adding an antibiotic perturbation reshapes this field; sweeping the most common antibiotic perturbations reveals category-specific flow patterns that shift the microbiota toward different states. Some antibiotics redirect the flow toward the *Enterococcus*-dominated region of the map (e.g., penicillin).

**Figure 5.**
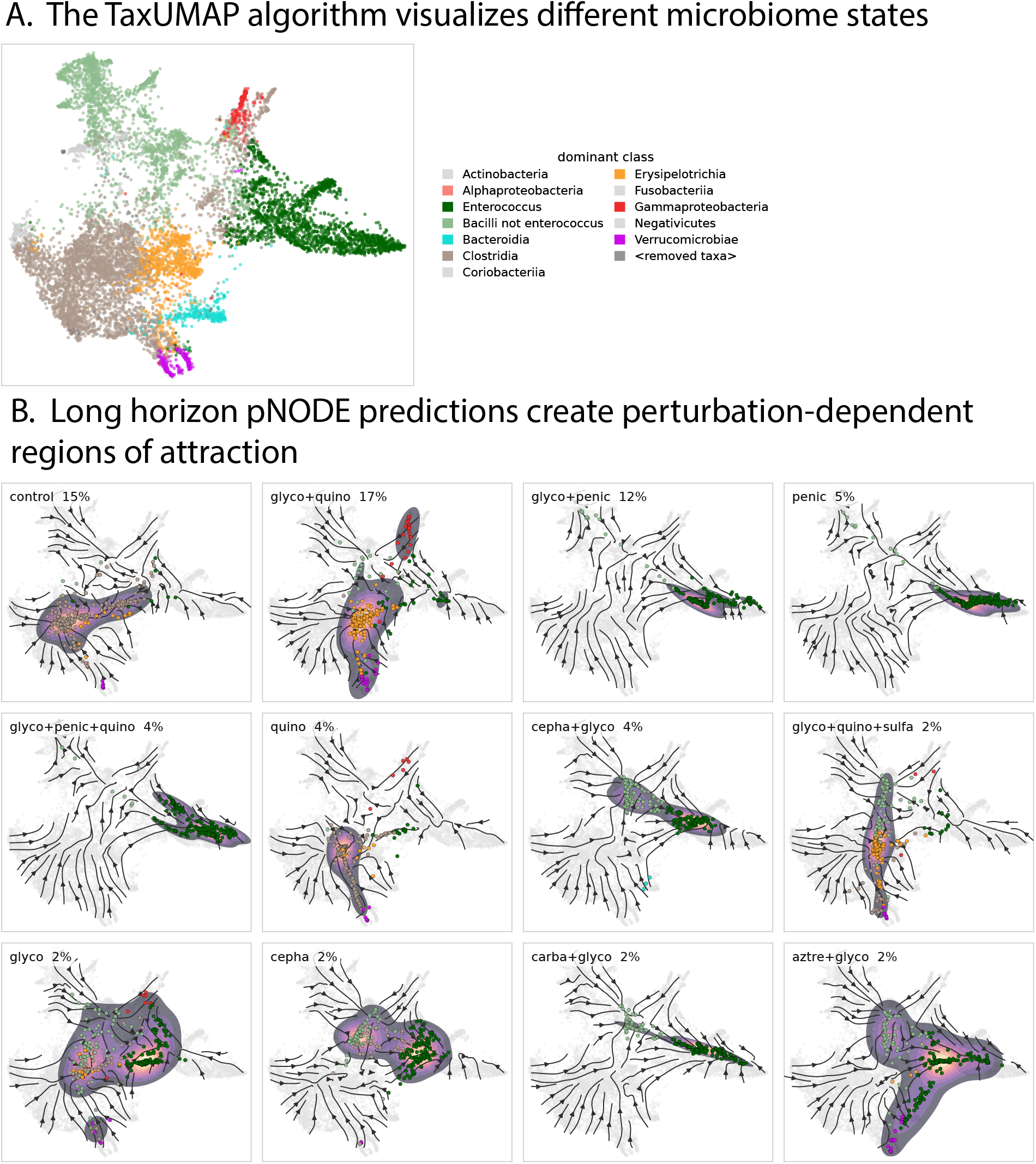
The pNODE encodes an antibiotic-dependent flow field over community-composition space. (A) Reference TaxUMAP map of the cohort, each point one fecal sample with published TaxUMAP coordinates (10,346 samples from 1,169 patients), coloured by its dominant bacterial class (key at right). (B) pNODE-induced flow under the 12 most frequent antibiotic perturbation states, no-antibiotic control first. Streamlines show the direction of community motion obtained by advancing 2,500 observed communities one 0.5-day step under each condition. The fields are overlaid with long-horizon endpoints: the density (magma) and dominant class (points) of 250 diverse pre-transplant communities integrated forward 30 days under each condition. Each panel is a perturbation *state*—the exact set of antibiotic categories active simultaneously rather than a single drug—because 68.2% of consecutive sample transitions in the training data occur with two or more categories active at once. The percentage on each panel is that state’s share of training transitions; the 12 shown account for 70.6%. Category abbreviations: glyco, glycopeptides (vancomycin); quino, quinolones (ciprofloxacin, levofloxacin); penic, penicillins (piperacillin–tazobactam, ampicillin); cepha, cephalosporins (cefepime, ceftazidime); carba, carbapenems (meropenem); sulfa, sulfonamides (trimethoprim–sulfamethoxazole); aztre, aztreonam. Long-horizon endpoints extrapolate beyond the training window and reflect the learned model’s attractors rather than directly observed dynamics (see Methods).

Integrating a diverse set of starting communities to a long horizon under each condition, the endpoints concentrate at different ecological states, colored by the dominant bacterial class (Fig. 5B). These attractors are a property of the learned model and, at long horizons, extrapolate beyond the 7-day training window (see *Methods*). Together, this view shows that the pNODE has learned a structured, perturbation-dependent flow over composition space whose antibiotic-driven compositional flux toward *Enterococcus* domination provides the signal that is then used to inform the BSI-prediction analysis.

## Discussion

Here we introduce pNODE as a perturbation-aware framework for learning microbiome dynamics from clinical time-series data. The central result is that a continuous-time neural dynamical model can capture microbiota responses to antibiotic exposure in a way that is both predictive and biologically meaningful. In synthetic benchmarks, pNODE outperformed gLV inference even when the data were generated by a gLV model. This highlights that under the sparse, noisy, irregular sampling conditions typical of clinical microbiome studies, the formally correct model may not be the most useful for prediction. pNODE models the ecological effects of treatment by learning nonlinear dynamics directly from observed trajectories and perturbation histories while avoiding a fixed interaction structure.

We illustrated how pNODE-generated trajectories can predict the incidence of bloodstream infections by *Enterococcus* in allo-HCT patients, a clinical complication well known to be preceded by intestinal dominations. A key strength of this approach is that the model is trained only on microbial abundances and antibiotic timelines, yet the learned trajectories contain signal about downstream clinical outcomes. In allogeneic HCT patients, *Enterococcus* bloodstream infections are often preceded by intestinal expansion of *Enterococcus*. The pNODE captures this ecological transition: when run in generative mode from an initial fecal sample and the patient-specific antibiotic record, it produces predicted *Enterococcus* trajectories whose time-averaged abundance predicts subsequent *Enterococcus* BSI. Because infection labels were never used during training, the predictive performance suggests the model learned aspects of microbiome dynamics upstream of infection risk rather than fitting the outcome.

The comparison between taxonomic resolutions also provides useful insight into how microbiome dynamics can be represented for prediction. The class-level model uses a compact state space that enables direct comparison with gLV on the qPCR-restricted subset, while explicitly resolving *Enterococcus* as a clinically important coordinate. The genus-level model preserves more biological detail and can be trained on the larger full cohort. These results show that pNODE can operate across taxonomic scales and that the choice of scale can balance interpretability, data availability, and predictive signal.

Beyond forecasting individual trajectories, projecting the genus-level pNODE into TaxUMAP space reveals antibiotic-dependent flow fields over community composition. This shows how antibiotics reshape the direction and magnitude of community dynamics across the microbiome landscape. Some perturbation states steer communities toward regions dominated by *Enterococcus*, whereas others steer the microbiota to different compositions. These flows summarize dynamics learned from thousands of clinical samples and generate testable hypotheses about how antibiotic regimens redirect communities toward clinically relevant ecological states.

Several limitations should be considered. First, the clinical analyses were performed in a single-center allo-HCT cohort, where antibiotic exposure, sampling practices, patient population, and infection-control procedures may differ from those at other institutions or in other cancer-treatment settings. External validation will be needed to determine how well the learned dynamics transfer across centers and clinical contexts. Second, the current perturbation encoding represents antibiotic exposure as binary category-level treatment intervals. This captures major treatment structure but does not incorporate dose, route, pharmacokinetics, fecal drug concentrations, or other concurrent perturbations such as diet, chemotherapy, immune suppression, bowel preparation, or microbiome-directed interventions. Third, most analyses use relative abundances, which may not fully capture changes in absolute microbial burden. Fourth, even the higher genus-level resolution still aggregates substantial strain-level diversity, including clinically important variation in antibiotic resistance, virulence, and colonization potential. Finally, although pNODE predictions are clinically informative, they should not be interpreted as causal estimates of the effect of changing a patient’s antibiotic regimen without prospective validation or explicit causal modeling.

These limitations point to several future directions. The framework could be extended to include richer perturbation features, absolute-abundance measurements, strain-resolved profiles, metabolomic readouts, and host clinical variables. It could also be adapted to prospective decision-support settings in which candidate antibiotic regimens are evaluated in silico for their predicted ecological consequences before treatment decisions are made. More broadly, pNODE illustrates how clinical microbiome datasets can be used not only to identify associations between microbiota states and outcomes, but also to learn dynamical rules that forecast how patient-specific ecosystems respond to treatment [11]. Such models may ultimately support microbiome-aware antibiotic stewardship: selecting therapies that treat infection risk while minimizing ecological disruption and preventing pathogen domination.

## Supplementary figure legends

**Supplementary Figure S1. Full pNODE / gLV MAE-ratio heatmaps across the (noise, density, training-size) grid of in silico experiments.** Three two-dimensional slices of the synthetic-benchmark grid are shown, each colour-coded by the mean held-out test MAE ratio (pNODE / gLV) over replicates. Darker red = lower ratio = pNODE wins by a larger margin; every cell in every slice is < 1.0, indicating that pNODE outperforms gLV under all conditions tested. (Top) Noise level sigma vs sampling density at fixed training-size = 100. (Middle) Noise level sigma vs number of training trajectories at fixed density = 3. (Bottom) Number of training trajectories vs sampling density at fixed noise = 0.1. The colour scale is shared across all three panels and spans the global cell range of the filtered grid (0.10-0.95).

**Supplementary Figure S2. Direction-of-change accuracy on held-out clinical data: fraction of pNODE and gLV predictions whose sign of Delta *Enterococcus* matches the observed sign.** For every held-out index sample and every follow-up sample at integer time gap *T* days ahead (rounded to the nearest integer), we compare the sign of the predicted change in *Enterococcus* relative abundance to the sign of the observed change. Pairs with |Delta| < 0.005 are excluded as ties. (Top) Per-day direction accuracy at exactly *T* days ahead, smoothed with a 3-day rolling window. Solid red: pNODE (mean across 20 replicates); dashed green: gLV; dotted grey: chance (0.5). Shaded bands are ± 1 SE of the binomial proportion. pNODE accuracy is consistently above chance and above the gLV baseline across the full 1-60-day horizon. (Bottom) Cumulative direction accuracy: fraction of all (index, follow-up) pairs with gap ≤ *T* days whose predicted sign of Delta-*Enterococcus* is correct, plotted against *T*. The blue shaded region (right axis) shows the cumulative number of non-tie pairs available at each horizon, which grows from ∼650 at *T* = 5 days to ∼3,500 at *T* = 60 days. pNODE’s cumulative accuracy stabilises near 0.7 from ∼30 days onward, materially higher than gLV’s ∼0.6.

## Notes

### Competing Interest Statement

The authors have declared no competing interest.

### Summary of Updates

Corrected mistakes in code, added section on 'flow plots'. Added genus level pNODE experiments.

