## Supplementary figures and images for "Perturbation-Aware Neural ODE (pNODE) Learns Microbiome Dynamics from Clinical Data and Predicts Gut-Borne Bloodstream Infections in Patients Receiving Cancer Treatment"

### Supplemental Figure S1

A. Full InSilico results across many training data conditions
pNODE wins everywhere

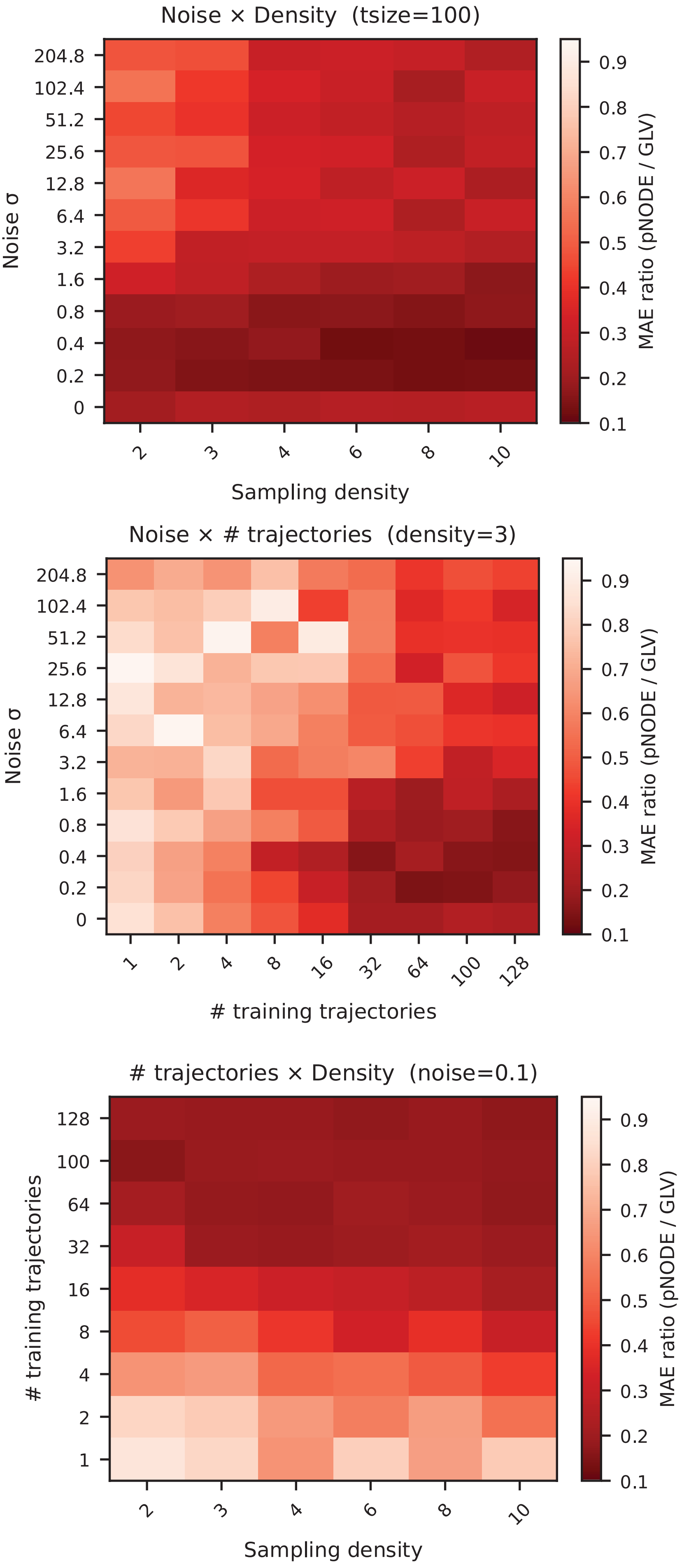

### Supplemental Figure S2

#### A. Direction accuracy of predicting sign of change of Enterococcus abundance

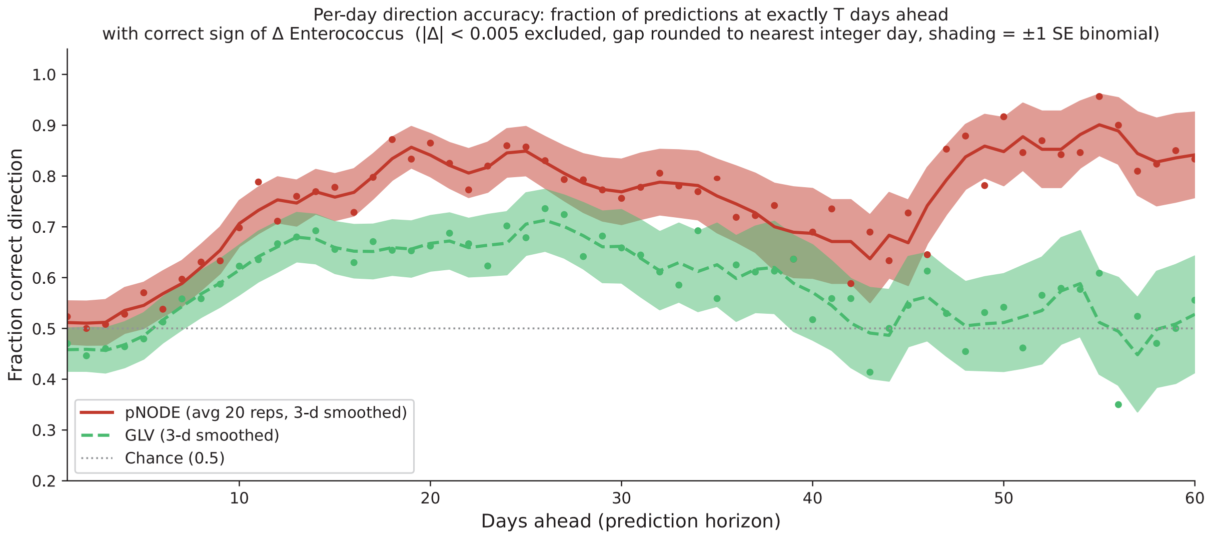


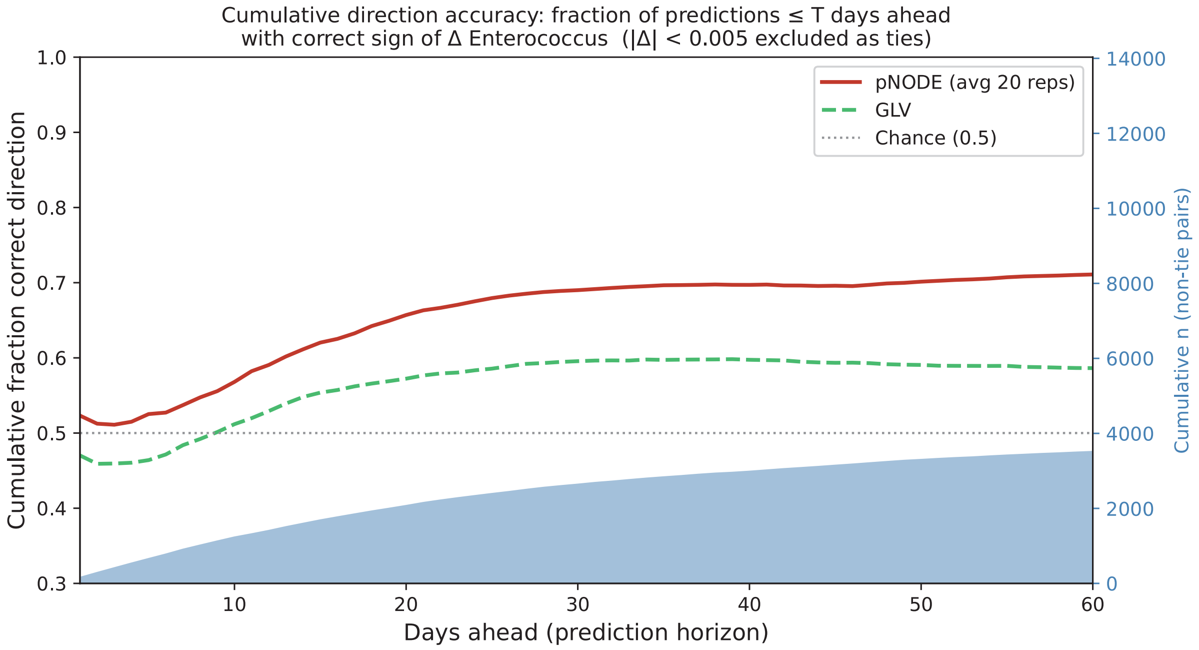
